# Learning Protein Structural Fingerprints under the Label-Free Supervision of Domain Knowledge

**DOI:** 10.1101/407106

**Authors:** Yaosen Min, Shang Liu, Chenyao Lou, Xuefeng Cui

**Affiliations:** Institute for Interdisciplinary Information Sciences (IIIS) Tsinghua University Beijing, China; E-Commerce Engineering with Law Beijing University of Posts and Telecommunications Beijing, China

## Abstract

**Finding homologous proteins is the indispensable first step in many protein biology studies. Thus, building highly efficient “search engines” for protein databases is a highly desired function in protein bioinformatics. As of August 2018, there are more than 140,000 protein structures in PDB, and this number is still increasing rapidly. Such a big number introduces a big challenge for scanning the whole structure database with high speeds and high sensitivities at the same time. Unfortunately, classic sequence alignment tools and pairwise structure alignment tools are either not sensitive enough to remote homologous proteins (with low sequence identities) or not fast enough for the task. Therefore, specifically designed computational methods are required for quickly scanning structure databases for homologous proteins**.

**Here, we propose a novel ContactLib-DNN method to quickly scan structure databases for homologous proteins. The core idea is to build structure fingerprints for proteins, and to perform alignment-free comparisons with the fingerprints. Specifically, the fingerprints are low-dimensional vectors representing the contact groups within the proteins. Notably, the Cartesian distance between two fingerprint vectors well matches the RMSD between the two corresponding contact groups. This is done by using RMSD as the domain knowledge to supervise the deep neural network learning. When comparing to existing methods, ContactLib-DNN achieves the highest average AUROC of 0.959. Moreover, the best candidate found by ContactLib-DNN has a probability of 70.0% to be a true positive. This is a significant improvement over 56.2%, the best result produced by existing methods**.

**GitHub: https://github.com/Chenyao2333/contactlib/**

**Index Terms:** **homologous proteins, protein structures, remote protein homolog detection, alignment-free comparisons**

## I. INTRODUCTION

Homologous proteins are the ones sharing a common ancestor. They carry critical information to understand protein functions and evolutions [1]. Thus, finding homologous proteins is usually the first and the indispensable step in many protein biological studies. Specifically, homologous proteins tend to share a common structure carrying a conserved function. By studying the correlation between the common structure and the conserved function, one could understand how these homologous proteins perform the conserved function. Understanding how proteins function is a critical step towards designing proteins with specific functions. This is the ultimate goal of current protein research.

As the number of known protein structures increases rapidly (i.e., more than 10,000 structures are deposited annually into PDB since 2016 [2]), the computational cost to perform a structure-based database scan for homologous proteins increases rapidly. One possible alternative solution is to apply the classic sequence alignment methods, such as BLAST [3]. Although such methods are highly efficient, their accuracies drop significantly for remote homologous proteins with low sequence identities [4], [5]. In such cases, structure similarities become the most reliable evidences for finding homologous proteins. This is mainly because structures are more conserved than sequences during the course of evolution. Therefore, it is highly desired to have an efficient and effective structure-based “search engine” for homologous proteins.

One intuitive way to find remote homologous proteins is to perform pairwise structure alignments between the query protein and all proteins in the database. In the past two decades, many successful methods have been developed for the pairwise structure alignment problem [6]–[14]. The main challenge here is to find the optimal superposition and the optimal residue matching simultaneously. Actually, this problem is known to be NP-hard [15], [16], and time-consuming heuristic algorithms are applied to find local optimal solutions. As a result, performing a database scan could take hours to days on a single computer [4], [5]. However, a “search engine” should return results in seconds, and thus existing pairwise structure alignment methods are not suitable for structure database scans.

In order to satisfy the speed requirement for quickly scanning structure databases, several successful methods have been proposed [17]–[19]. For example, FragBag [20] adopts the classic bag-of-word approach for the literature search problem, and treats protein structures as literatures with a bag of structure fragments. ContactLib [21] abstracts all contact groups within the protein structure, and represents them as low-dimensional fingerprint vectors for highly efficient indexing. SmotifCOMP [22] employs structure motifs at secondary structure element boundaries to significantly reduce the dimensionality of structure representations. Due to the tradeoff between speeds and accuracies, all these pioneer attempts achieve high speeds with noticeable losses on accuracies. Therefore, there is still room for accuracy improvements, and achieving higher accuracies with high speeds is the main challenge to scan a database for remote homologous proteins.

Deep neural networks (DNNs) have outperformed state-of-the-art methods in different research areas [23], including protein computational biology [24]. For examples, DNNs have been successfully used for solvent accessibility and secondary structure predictions [25], [26], contact map predictions [27], [28] and fold recognitions [29], [30]. One thing in common among these DNNs is that they all employ supervised learning, and thus training such DNNs usually requires a sufficiently large amount of labeled data. Recently, it has been shown that label-free supervision is possible, and domain knowledge is sufficient to train DNNs [31]. Specifically, DNNs have been successfully trained to track free falling objects in videos without any labeled data. The only training supervision used here is the physical law of free falling objects. Although such label-free supervision greatly eases the labeled data requirements of training DNNs, it has not yet been applied in protein computational biology.

In this manuscript, we propose a new method, called ContactLib-DNN, to perform fast database scans for remote homologous proteins in the following steps: first, contact groups are abstracted from the query protein structure; second, a DNN is employed to convert contact groups into low-dimensional vectors as fingerprints (see Fig. 1 for an illustration of the first two steps); third, these fingerprints are used to scan the database for similar contact groups (with pre-computed fingerprints) using a highly efficient indexing algorithm; fourth, using these similar contact groups, an alignment-free similarity score is computed between the query protein and each protein in the database; and finally, the proteins in the database are ranked according to the similarity scores.

**Fig. 1:**
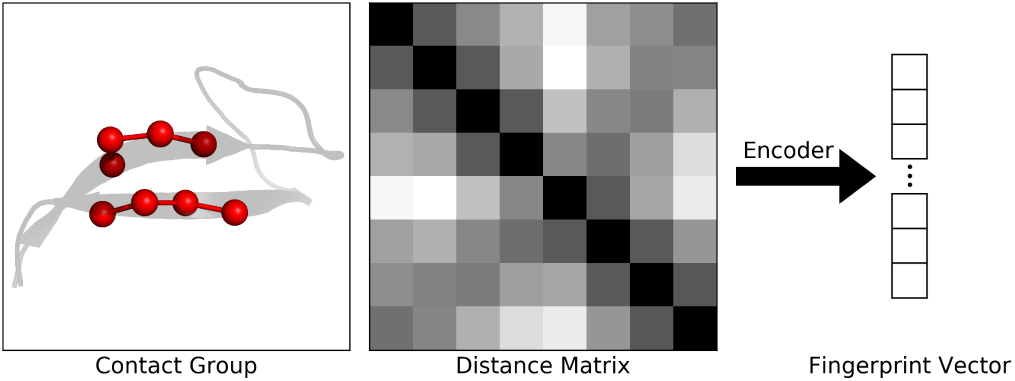
Conducting vector representations of contact groups: the 3-dimensional structure (left) of a contact group (red) is first converted to a residue-residue distance matrix (middle); the distance matrix is then converted to a low-dimensional vector (right) by a deep encoder (arrow, see Fig. 2); and the vector is finally used as a fingerprint to represent the contact group.

Our new ContactLib-DNN method improves the original ContactLib [21] method in two aspects: (1) the revised definition of contact groups better eliminates noisy (i.e., useless) and redundant contact information; and (2) the DNN converts contact groups into fingerprint vectors such that *the Cartesian distance between two fingerprint vectors well matches the RMSD between the two corresponding contact group structures*, as shown in Fig. 2. Here, RMSD serves as the domain knowledge to supervise our training process. As a result, fingerprint vectors with lower dimensionalities are conducted for higher speeds, and more precise indexing results are achieved for higher accuracies. To the best of our knowledge, ContactLib-DNN is the first DNN trained under label-free supervision of domain knowledge in protein computational biology.

**Fig. 2:**
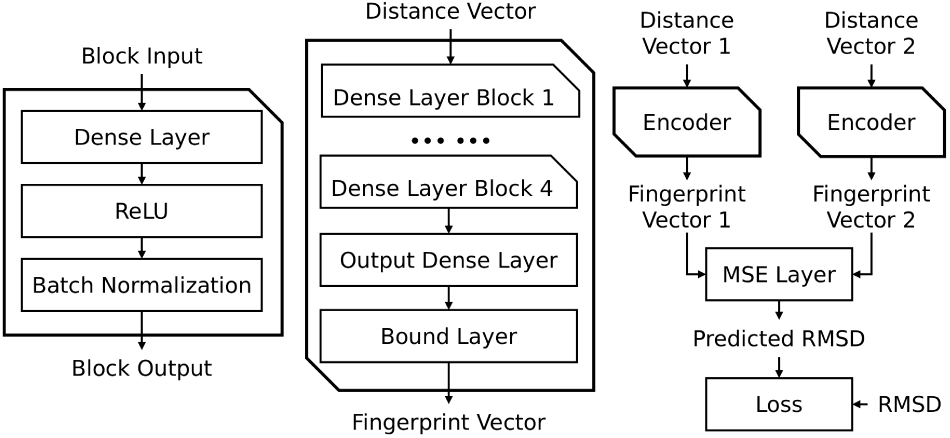
The architecture of the deep encoder: a dense layer block (left) is built to learn the hidden features; an encoder (middle) is built to generate the fingerprint vectors representing the contact groups; the number of neurons of the pre-activation output dense layer equals to the dimension of the fingerprint vector; the bound activation layer trim the outputs into range [-8Å, 8Å]; and the objective of the training process (right) is finding a fingerprint vector space such that the mean square error (MSE) between two fingerprint vectors equals to the RMSD between the two corresponding contact groups.

The performance of our ContactLib-DNN is compared to state-of-the-art methods, FragBag [20] and ContactLib [21]. It has been observed that the AUROC distribution of ContactLib-DNN is much closer to the maximum value of one with the highest average AUROC of 0.959. When focusing on the top ten candidates found by ContactLib-DNN, ContactLib and FragBag, the precisions are 0.427, 0.316 and 0.019, respectively. Therefore, all results support that ContactLib-DNN is the most accurate method.

## II. METHOD

### A. Contact Group Definition

It has been reported that one contact for every twelve residues allows accurate protein structure modeling [32]. Thus, residue-residue contacts are capable of determining protein structures, and we introduce contact groups to capture the critical contacts and their environments for finding structurally similar proteins.

A contact group is defined as a pair of contacting fragments with the same length of *l* (i.e., the number of residues of the fragment). Fig. 1 shows one contact group with *l* = 4 on the left. Here, each residue is represented by its C_*α*_ atom, and *l* can be used to control how much sequential environment to be included. The C_*α*_-C_*α*_ distance threshold of *d*_*c*_ is used to approximately determine if two residues are in contact. Moreover, a contact group should contain at least *n*_*c*_ contacts between the two fragments to effectively exclude noises introduced by our contact approximation. Finally, the maximum pairwise C_*α*_-C_*α*_ distance of *d*_*g*_ is introduced to control how much spacial environment to be included.

Interestingly, we observed that different contact groups may differ in capability on modelling structures. Thus, a filtering step is performed based on the secondary structures for higher accuracies and higher speeds.

### B. RMSD-Supervised Deep Encoder

The motivation to use a deep encoder lies in the redundant representation of contact groups and the RMSD restraint. Specifically, taking *i* as the sequence number defined in the PDB format, the average distance between two *α*-helix residues *i* and (*i* + 4) is 6.22Å with a standard deviation of 0.34Å. Such a low standard deviation implies a high redundancy. As illustrated in Fig. 1, the distance matrix of a contact group is encoded into a fingerprint vector to eliminate such redundancies. Moreover, traditional methods are not designed to learn fingerprint vectors under the RMSD restraint. For examples, PCA [33] can only conduct a linear compression; t-SNE [34] is merely able to yield two or three dimensional vectors; word2vec [35] is more suitable for extracting between 500 and 1,000 dimensional information. Therefore, the specifically designed deep encoder in Fig. 2 is introduced to produce compact fingerprint vectors under the RMSD restraint.

The deep encoder is shown in the middle of Fig. 2, and it is used to produce fingerprint vectors for contact groups. Since we do not have labels for such fingerprint vectors, a supervised learning is not possible. Thus, we introduce the Y-shaped RMSD-supervised learning model to train the encoder, as shown on the right of Fig. 2. Specifically, the two encoders in the Y-shaped model share the parameters, and hence only one encoder is trained. The learning objective is the RMSD restraint such that the MSE between a pair of fingerprint vectors should equal the RMSD between the two corresponding contact groups. Therefore, RMSD serves as the domain knowledge to supervise the label-free learning process.

### C. Searching the Library of Contact Groups

Before introducing the searching algorithm, we first describe the evaluation of the similarity between two fingerprint vectors. For two fingerprint vectors *V*_*q*_ and *V*_*t*_ from the query protein and a target protein, the distance between *V*_*q*_ and *V*_*t*_ is determined by the *l*_∞_-norm. Then, *V*_*q*_ and *V*_*t*_ are considered to be similar by the following constraint:

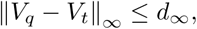

where *d*_*∞*_ is a user-defined threshold. Then, an indexing algorithm can be used to quickly retrieve all similar contact groups in the target database (see [21]). In order to use the indexing algorithm, real values are discretized into multiple bins. For example, one bin could represent 0.1 Å, and hence *d*_*∞*_ = 1.2Å is equivalent to 12 bins.

To retrieve homologous proteins given a query protein, all contact groups of the query protein are first abstracted and transformed to fingerprint vectors. Then, the indexing algorithm with the *l*_*∞*_-norm distance function and an user-specified *d*_*∞*_ threshold is used to find all pairs of similar contact groups between the query protein and any target protein in the target database. The number of similar pairs (i.e., hits) between the query protein and a target protein can be simply counted to calculate the similarity score defined as follows:

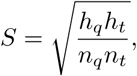

where *h*_*q*_ and *h*_*t*_ are the number of hits on the query protein and the target protein, respectively; and *n*_*q*_ and *n*_*t*_ are the number of contact groups of the query protein and the target protein, respectively. Thus, higher scores imply higher similarities, and all target proteins in the target database are ranked according to the similarity score for output.

## III. RESULTS

### A. CULLPDB25 Dataset

In order to evaluate the performance of our method, a dataset of non-redundant and high-quality protein structures is prepared with the help of the widely used PISCES server [36]. Specifically, the sequence identity is less than 25%, the resolution is less than 2.0Å and the R-factor is less than 0.25. This protein list was downloaded on Jan. 15, 2018, and then only proteins with more than 50 residues and less than 1,000 residues were kept. As a result, 8,437 proteins were collected, and this dataset will be referred as CULLPDB25 in this manuscript.

### B. Model Settings

In our definition of contact groups (described in Section II-A), there are four hyperparameters, and the actual values used in our experiments are as follows: the minimal number of contacts *n*_*c*_ = 2, the fragment length *l* = 4, the maximum contact distance *d*_*c*_ = 8.0Å, and the maximum group distance *d*_*g*_ = 16.0Å. Our results also suggest to use RMSD cutoffs between 0.6Å and 1.4Å. Finally, contacts groups containing only *α*-helix and coil fragments are filtered out for higher speeds and higher accuracies. For the sake of saving space, the details of yielding above model settings are eliminated.

### C. Encoding Fingerprint Vectors

Recall that the objective of our DNN training process is finding a fingerprint vector space such that the MSE between two fingerprint vectors equals to the RMSD between the two corresponding contact groups. In this section, we first demonstrate that the DNN outperforms the classic PCA [33] in terms of our objective. Then, we demonstrate the positive impacts of our training objective on finding similar contact groups.

In this experiment, 100 million contact group pairs were randomly selected from CULLPDB25 as our training dataset, and 100 thousand contact group pairs were randomly selected as our testing dataset. We used the training dataset to train a deep encoder with 7 output neurons (described in Section II-B), and the training process was stopped after 100 epochs. Here, neither the optimal number of output neurons nor the optimal training process was the focus of this study. Thus, this setting was used for all the following experiments. The following analysis was conducted with the testing dataset and the DNN model after 100 epochs.

The heatmaps in Fig. 3 show the relationships between the MSE and the RMSD using fingerprint vectors encoded by DNN and PCA. It can be observed that the MSE of DNN encoded vectors is tightly close to the RMSD, whereas this is not the case for PCA encoded vectors. If we use the *l*_*∞*_ norm (also used by our indexing algorithm as described in Section II-C) as the distance metric between fingerprint vectors to predict similar contact groups, the prediction accuracies are shown in Table I. It can be observed that PCA tends to have more balanced values between precisions and recalls. On the other hand, DNN is capable of delivering near-optimal precisions (e.g, 0.985) while discovering majority of similar contact groups (i.e., recall > 0.5). This is a desired phenomenon caused by our DNN training objective, and this phenomenon leads to a better overall performance in the next section.

**Fig. 3:**
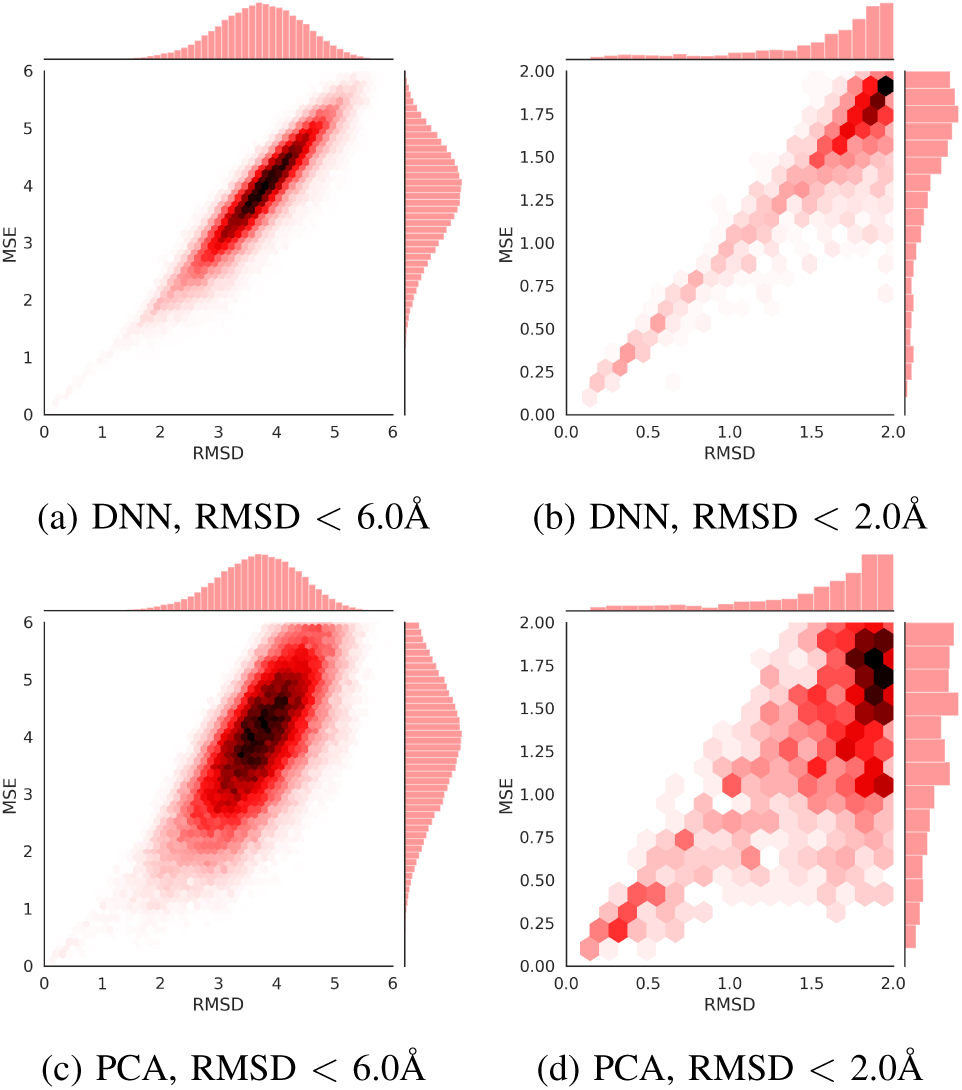
Comparisons of the fingerprint vectors conducted by DNN and PCA: recall that for high-quality fingerprint vectors, the MSE between two fingerprint vectors should well match the RMSD between the two corresponding contact groups; in general cases, the heatmap of DNN (a) shows a narrower distribution along the diagonal comparing to that of PCA (c); especially, when focusing on the similar contact groups with RMSD < 1.5, the heatmap of DNN (b) shows a solid relationship between MSEs and RMSDs, while this is not the case for PCA (d); and comparing to PCA, the fingerprint vectors conducted by DNN have an outstanding capability to distinguish similar contact groups from dissimilar ones.

**TABLE I:** Accuracies to Find Similar Contact Groups

| RMSD Cutoff |  | 0.6 | 0.8 | 1.0 | 1.2 | 1.4 |
| --- | --- | --- | --- | --- | --- | --- |
| Precision | PCA | 0.839 | 0.860 | 0.712 | 0.695 | 0.617 |
|  | DNN | **0.943** | **0.985** | **0.978** | **0.950** | **0.973** |
| Recall | PCA | **0.584** | **0.688** | **0.646** | **0.596** | **0.503** |
|  | DNN | 0.371 | 0.536 | 0.540 | 0.500 | 0.451 |

### D. Overall Performance Evaluations

In this experiment, the overall performance of our new ContactLib-DNN to scan structure databases for homologous proteins is evaluated. Then, the results are compared to Frag-Bag [20], ContactLib [21] and ContactLib-PCA, a variant of ContactLib-DNN using a PCA in place of the DNN. Since neither the source code nor the Smotif library is publicly available for SmotifCOMP [22], it is not available for comparisons in this experiment.

For each protein in CULLPDB25, we used it as the query protein and the rest of CULLPDB25 as the target database (i.e., leave-one-out cross validation). Then, ContactLib-DNN was used to scan the target database for homologous proteins. Specifically, for each protein, the contact groups were abstracted based on our hyperparameters. For each contact group, a 7-dimensional fingerprint vector was generated using our deep encoder (see Section III-C). Then, we used the contact groups of the query protein and the contact groups in the target database to find homologous proteins (see Section II-C). Here, two proteins are presumed to be homologous if their TM-score is greater than 0.5. Recall that there are 8,437 proteins in CULLPDB25. Interestingly, there are 1,176 proteins without any homolog. Since AUROCs are not defined for such cases, we used the remaining 7,261 proteins in the following analysis.

AUROC [37] is capable of describing the sorting capability of a search tool. If a positive and a negative sample are randomly selected, AUROC evaluates the probability that the positive sample ranks prior to the negative one. Fig. 4a shows the average AUROC among the 7,261 protein structures. It can be seen that ContactLib-DNN’s average AUROC achieves 0.959, which is improved by 1.9%, 4.5% and 27.9% compared to ContactLib-PCA, ContactLib and FragBag, respectively. Furthermore, from the AUROC distributions in Fig. 4b, 75% of ContactLib-DNN’s AUROC results exceed 0.945, whereas only around 49% of ContactLib’s AUROCs reach the same value. Therefore, ContactLib-DNN has evident advantages on AUROCs, which is sufficient to verify the sorting effectiveness of a search tool.

**Fig. 4:**
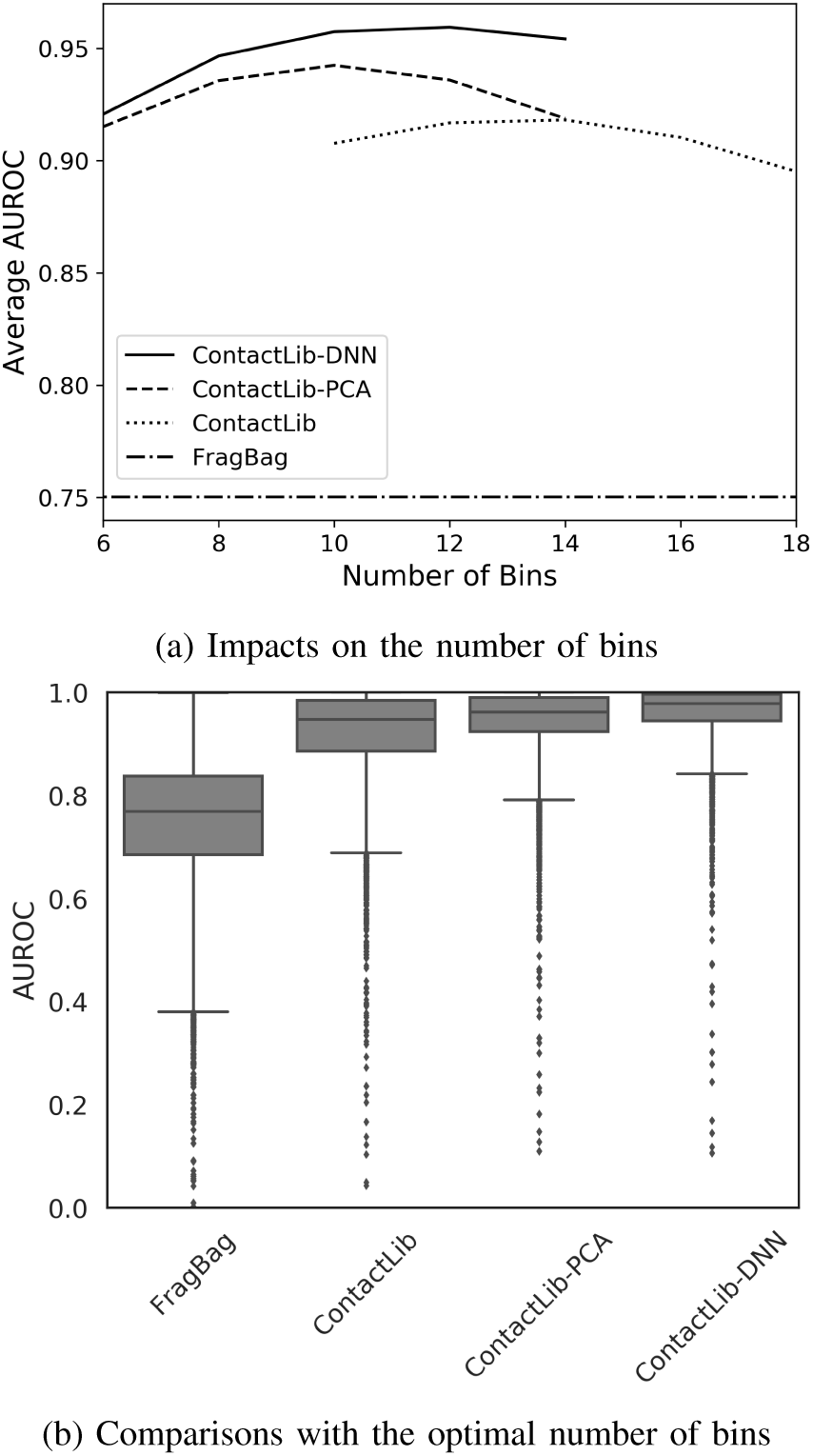
Evaluations of scanning structure databases for homologous proteins: every protein in the CULLPDB25 dataset is used to find homologous proteins from the rest of the dataset; an AUROC is calculated for each database scan; the number of bins is an user-specified threshold equivalent to the *l*_*∞*_-norm distance threshold *d*_*∞*_ (see Section II-C); (a) the relationships between the average AUROC and the user specified number of bins are plotted; (b) the AUROC distributions with the optimal number of bins are shown in the box-and-whisker plot (the whisker height is twice of the box height); and comparing to other tested methods, ContactLib-DNN achieves the highest average AUROC of 0.959 with AUROC distributions much closer to the perfect value of one.

The precision of top predictions is another index to evaluate the effectiveness of search engines. Specifically, the principle for a search engine is listing as many true positive results as possible on the top of the list, which can be evaluated by the precision of top predictions. Table IIa shows the precision of top 1, 10, 20, and 40 predictions. Please note that the average number of positives is approximately 34. Comparing to ContactLib-PCA, ContactLib-DNN promotes the average precision significantly by 25.6%. In addition, we calculate the probability of precision > 0.0 and precision > 0.5. The probability of precision > 0.0 indicates the likelihood that at least one true positive result emerge in top predictions. According to Table IIb, the probability of having at least one true positive in top 10 predictions by ContactLib-DNN reaches 0.85, whereas the other methods reaches at most 0.77. The probability of precision > 0.5 indicates the likelihood that the majority of the top predictions are true positives, and ContactLib-DNN outperforms the others significantly as well.

**TABLE II:** Evaluations of Top Predictions

(a) Average precisions
| Method | Top 1 | Top 10 | Top 20 | Top 40 |
| --- | --- | --- | --- | --- |
| FragBag | 0.035 | 0.019 | 0.017 | 0.017 |
| ContactLib | 0.562 | 0.316 | 0.247 | 0.184 |
| ContactLib-PCA | 0.583 | 0.340 | 0.269 | 0.203 |
| ContactLib-DNN | **0.700** | **0.427** | **0.340** | **0.257** |

(b) Probability of precision > 0.0
| Method | Top 10 | Top 20 | Top 40 |
| --- | --- | --- | --- |
| FragBag | 0.172 | 0.251 | 0.351 |
| ContactLib | 0.758 | 0.804 | 0.852 |
| ContactLib-PCA | 0.767 | 0.817 | 0.865 |
| ContactLib-DNN | **0.850** | **0.883** | **0.913** |

(c) Probability of precision > 0.5
| Method | Top 10 | Top 20 | Top 40 |
| --- | --- | --- | --- |
| FragBag | 1.4e-4 | 0 | 0 |
| ContactLib | 0.266 | 0.182 | 0.102 |
| ContactLib-PCA | 0.294 | 0.208 | 0.126 |
| ContactLib-DNN | **0.394** | **0.291** | **0.198** |

Although AUPRC is widely used to evaluate prediction models, it is not suitable in the context of search engines. This is mainly because, for a search engine, its users usually focus on the precision of top predictions rather than the overall accuracies. Nevertheless, the AUPRC for ContactLib-DNN reaches 0.433, which is 35.3% and 50.9% higher than ContactLib-PCA and ContactLib, respectively. Thus, the observations of AUPRC are consistent with those of AUROC.

## IV. CONCLUSIONS

In this manuscript, we propose a novel method, ContactLib-DNN, to rapidly scan structure databases for homologous proteins. The superiority of ContactLib-DNN is reflected in both efficiency and effectiveness. With respect to the efficiency, the contact redundancy elimination, the DNN dimension reduction and the bitwise indexing significantly reduces the search time. As a result, the average CPU time to scan the CULLPDB25 dataset for a query structure is merely 1.7 seconds using a single core of an Intel(R) Core(TM) i7-8700K 3.70GHz processor. Moreover, our program requires less than 2GB memory.

The improvement of ContactLib-DNN’s effectiveness results from two core ideas. First, filtering out the three most redundant contact group classes evidently improves the AU-ROC and the AUPRC while reduces the runtime complexity. Second, ContactLib-DNN compresses contact groups into fingerprint vectors, whose Cartesian distances well matches the RMSDs between corresponding contact groups. This key training constraint achieves a near-perfect precision of 0.985 while discovering the majority of similar contact groups, as shown in Table I. These two core ideas lead an improvement of AUROC to 0.959 in the overall experiment. In summary, ContactLib-DNN is the most accurate method among ContactLib-DNN, ContactLib-PCA, ContactLib [21] and FragBag [20].

